# Environment dependent benefits of sociality in Soay sheep

**DOI:** 10.1101/2025.05.04.652099

**Authors:** Erin R Siracusa, Xavier Bal, Delphine De Moor, Gregory F Albery, Ross Beattie, Sanjana Ravindran, Ellis Wiersma, Amelie Renwick Wilson, Josephine M Pemberton, Daniel H Nussey, Matthew J Silk

**Affiliations:** School of Psychology, Centre for Research in Animal Behaviour, University of Exeter, Exeter, UK; Institute of Ecology and Evolution, School of Biological Sciences, University of Edinburgh, Edinburgh, UK; Department of Primate Behavior and Evolution, Max Planck Institute for Evolutionary Anthropology, Leipzig, Germany; School of Natural Sciences, Trinity College Dublin, Dublin, Ireland

**Author notes:** corresponding authors: ERS -; MJS.

**Keywords:** environmental variation, fission-fusion, fitness, mammals, relationship quality, relationship quantity, social networks, social buffering, survival

## Abstract

An individual’s social connections have strong effects on fitness. Despite this, there is pronounced among-individual variation in social behaviour. This variation may be maintained if different types of social connections have environment-dependent fitness benefits, but this has rarely been tested. We applied network analysis to 37 years of social association data in Soay sheep to test this hypothesis. Our results show that both relationship quantity (having many association partners) and relationship quality (having strong associations) are linked with survival. Crucially, the relative importance of relationship quantity and quality depends on overwinter weather, with harsh conditions favouring quantity over quality, likely due to thermoregulatory benefits. Relationship quality, meanwhile, was most beneficial under benign weather conditions, possibly due to enhanced familiarity with conspecifics. These findings advance our understanding of social benefits in fission-fusion societies and show that changing environmental conditions may be an important mechanism maintaining variation in social behaviour in natural populations.

## Introduction

The social structure of animal populations has important ecological and evolutionary consequences (Kurvers et al. 2014), for example by shaping the spread of infectious diseases (Bansal et al. 2007; Silk & Fefferman 2021) or determining who can cooperate with whom (Brask et al. 2019; Fehl et al. 2011). Social structure emerges from the patterns of social interactions that occur among members of the population, which themselves also shape individuals’ ability to access social information (Aplin et al. 2012; Snijders et al. 2021), acquire food (Ellis et al. 2017; Marshall et al. 2015; Snijders et al. 2021), find mates (Oh & Badyaev 2010), and care for young (Weidt et al. 2008). As a consequence, various measures of social connectedness have been shown to be consistently associated with health, survival and reproductive success across a wide range of taxa (Formica et al. 2012; McDonald 2007; Riehl & Strong 2018; Snyder-Mackler et al. 2020). However, despite these strong associations with fitness-related traits there is often considerable variation among individuals in their social networks across multiple axes (Bergman & Beehner 2015; Dakin & Ryder 2020; Ellis et al. 2019; Kutsukake 2009; Lukas & Clutton-Brock 2018; Schülke et al. 2022). Some individuals may have many social partners while others focus their social interactions on a few particular partners. At a network level, some individuals may be much more integrated within groups or cliques, while others may act as important bridges between different parts of the population. Quantifying the factors shaping the social interactions individuals engage in, which scale up to structure their social networks, is therefore critical to revealing why the social systems of different populations and species vary.

One potential explanation for the observed variation in social behaviour is that the adaptive benefits of social interactions are context-dependent (Bell 2010; Brown et al. 2016; Chevin & Haller 2014; Formica et al. 2021; Guindre-Parker & Rubenstein 2020; Testard et al. 2024; Wice & Saltz 2021). Social networks often change seasonally and/or in response to changing resource distributions (Bonnell et al. 2023; Foster et al. 2012; Henzi et al. 2009; Kusch & Lane 2021; Nandini et al. 2017; Prehn et al. 2019; Silk et al. 2018), and the benefits of occupying particular social network positions can also vary according to environmental conditions (Ellis et al. 2017; Fisher & Cheney 2023). For instance, social interactions can buffer or shield individuals against the consequences of harsh environmental conditions (Cornwallis et al. 2017; Fisher et al. 2021; Guindre-Parker & Rubenstein 2018; Kingma 2017; McFarland et al. 2017; McFarland & Majolo 2013; Testard et al. 2024; Warrington et al. 2024). For example, in killer whales (Orcinus orca) males who are more socially integrated are more likely to receive food from other group members, which reduces mortality risk, especially in years of low prey abundance (Ellis et al. 2017). These results suggest that different environmental pressures require distinct social solutions, which will cause the adaptive value of social relationships to vary considerably over time and space (De Moor et al. preprint). Understanding the extent to which and how the environment shapes selection on social behaviour offers an important window into the mechanisms underpinning the effects of sociality on fitness, while also helping us to understand the processes that maintain among-individual variation in social network position. Yet, to date there is very little research on the context dependent benefits of sociality.

A major axis of variation along which social relationships vary is in the formation of many weaker ties, which widely connects individuals to their networks as a whole (i.e. relationship ‘quantity’), versus the formation of a few stable, stronger relationships with key partners (i.e. relationship ‘quality’) (De Moor et al. preprint; Ellis et al. 2019; Schülke et al. 2022). Both quantity and quality of social ties have been empirically linked to fitness proxies (Carter et al. 2017; Gerber et al. 2022; Ostner & Schülke 2018). Moreover, both dimensions tend to be moderately or negatively correlated (although evidence for this is taxonomically limited; (Schülke et al. 2022; de Silva et al. 2011), suggesting they capture fundamentally different aspects of sociality (De Moor et al. preprint), and might even trade off. When these different social strategies are likely to be selected for is highly environment-dependent as they typically provide distinct benefits. Relationship quantity is hypothesized to be favoured under circumstances where social benefits are a numbers game - and are group-dependent rather than relationship-based (De Moor et al. preprint). For example, under cold or wet weather conditions individuals stand to gain important thermoregulatory benefits by huddling with many partners for warmth (Campbell et al. 2018; Paquet et al. 2016). Relationship quality, on the other hand, should be important when navigating competition over valuable and defendable resources like mates, territories, and food (De Moor et al. preprint). In such cases, having strong and consistent social relationships can provide benefits such as reduced aggression and social vigilance (Favreau et al. 2015; Fox et al. 2024; Griffiths et al. 2004; Siracusa et al. 2019) and/or increased cooperation to acquire and/or protect crucial resources (Connor et al. 2001; Dale et al. 2017; Heesen et al. 2014; Kern & Radford 2021). Consequently, if environmental conditions vary over time and different conditions favour either the quantity or quality of social relationships, then this will naturally help to maintain variation in social behaviour within a population. However, this idea has yet to be tested in wild populations.

Most work on the overall and context-dependent fitness benefits of sociality remains limited to species in stable, complex societies (Ostner & Schülke 2018; Snyder-Mackler et al. 2020). As a result, we know little about the value of social relationships in the many species with more fluid patterns of social associations (but see Carter et al. 2009; Menz et al. 2017; Montero et al. 2020; Prehn et al. 2019; Vander Wal et al. 2015; Wolf et al. 2018). Fission-fusion social systems in which animals aggregate in groups, but membership of these groups varies considerably over time and space are widespread in nature (Aplin et al. 2012; Croft et al. 2003; Fortin et al. 2009; Kerth et al. 2006). Individuals in these societies enjoy various advantages of sociality such as reduced predation risk (Isvaran 2007; Kelley et al. 2011) or improved access to social information (Aplin et al. 2012), but with greater flexibility in who they associate with and when. In such systems we would expect social relationships to be less well established on average, and for individuals to interact with a wider social network (Silk et al. 2014). Consequently, the benefits of sociality might be very different in these fluid social systems and might vary more strongly across contexts, compared to more stable, complex societies. Therefore, quantifying the fitness benefits of sociality in a broader suite of social systems, and their variation across time and context, represents an important challenge for our understanding of the adaptive function of social behaviour. This could be key to unpacking the role of environmental variation in maintaining social variation.

Here we address this question by studying the social association patterns of adult female Soay sheep (*Ovis aries*) on the island of Hirta, St Kilda. The sheep resident in Hirta’s Village Bay area have been individually marked and monitored since 1985, providing high quality demographic data across multiple generations and over a range of environmental conditions (Clutton-Brock & Pemberton 2004). Soay sheep have a fluid, fission-fusion social system in which individuals typically feed and rest in groups but membership of these groups is dynamic. Most Soay sheep mortality occurs during late winter (February-March), due to interacting pressures of food limitation, winter weather and infection with gastro-intestinal parasites. Mortality is highest following a) years of high sheep numbers, due to high resource competition and reduced plant biomass, and b) wet, stormy winters, which creates thermoregulatory challenges for the sheep (Coulson et al. 2001).

Together, this creates a perfect natural experiment to test if social relationships help individuals cope with environmental challenges in fission-fusion societies, and whether the types of relationships that aid coping differ across varying environmental contexts. We focus on two broad aspects of an individual’s social network: i) relationship quantity (the number and distribution of an individual’s social connections) and ii) relationship quality (the extent to which individuals repeatedly associate with the same social partners). Broadly we predict that both relationship quantity and quality will enhance survival. However, we expect that some of those benefits will be context-dependent. Specifically, we predict that relationship quantity will be particularly important in enhancing survival prospects during challenging winters through enhancing huddling behaviour and access to shelter which are both thought to be important ways to enhance thermoregulation in sheep. In contrast, we expect that both quantity and quality will be important when access to food impacts survival. Specifically, we predict that having many social associates will improve social tolerance around limited (but widely dispersed) resources, while having strong overall connections to social partners will enhance familiarity and predictability of partners, reducing time spent in conflict. Our full set of social variables and associated predictions are provided in Table S1.

## Methods

### Data collection and study system

The population of Soay sheep (Ovis aries) in Village Bay, Hirta – an island in the remote St Kilda Archipelago, Scotland [57°49’N, 8°34’W] - has been subject to individual-based monitoring since 1985 (Clutton-Brock & Pemberton 2004). This is a free-living population with no predators and no interspecific competition for food resources, and which is largely free from human influence (Catchpole et al. 2000). Each spring, lambs are caught and given uniquely numbered ear tags within a few days of birth and are subsequently followed throughout their lives.

Each year, the population is censused approximately 30 times (range 20-38) over the course of three field expeditions: ten censuses each in spring (March–April), summer (July–August), and autumn (October–November), totaling 1098 censuses in total between 1986 and 2023. During censusing, observers walk three fixed routes simultaneously and record the location of all individuals to the nearest 100 m, as well as group membership. Group membership is assigned by experienced field workers using a chain-rule, where all individuals within 10m of another individual are deemed to be associating (mean 64 groups observed per census, range 3-150) (Albery et al. 2022; Castles et al. 2014).

The vast majority of mortality in this population occurs in late winter (85% of adult deaths occur in January -April; Clutton-Brock & Pemberton 2004), and daily carcass searches are carried out during field trips early in each year. Regular censusing, in combination with these mortality searches, means that an individual’s disappearance from the population or death date are known with a high degree of accuracy (Clutton-Brock & Pemberton 2004). Using this information, we could determine overwinter survival, which we measured as the probability of surviving until May 1^st^ of the following year.

### Social network construction

We built annual social association networks for the period 1986-2023 using group membership data from the censuses for adult females only (> 2 years old; 4550 female-years over 968 females). We focused on female-female associations to estimate the effects of social associations on survival in isolation of potential socio-sexual influences that might be captured by including data from both sexes. Typically, individuals were only recorded the first time they were seen on a given census route, despite the fact that they might have been observed again in another group later in the census. While this has the potential to lead to missed observations of associations, and could lead to biases, information from observers that this happens relatively infrequently, combined with a simulation study demonstrate it is unlikely to substantially affect the networks constructed or conclusions drawn about individual social network position (see Supplementary Materials). On the occasions where an individual was recorded more than once, either on the same census route or on different census routes, we retained any observations of the same individual that were ≥60 min apart to help maximize the use of available information when generating our networks, while avoiding excluding observations that were likely to represent non-independent observations of the same group. To confirm that this choice did not influence our results, we replicated our analyses while retaining only the first observation for each individual from a given census and found similar results (see Fig. S1).

In addition, individuals varied in the unique number of censuses they were observed in each year (mean: 20, range: 1 – 34). Reduced sampling of networks leads to greater uncertainty in the true social relationships between individuals (De Moor et al. 2024; Hart et al. 2023) and the position of individuals in the network (Silk et al. 2015). Further, or perhaps as a consequence of this, there was an association between the number of census observations and the individual social metrics we estimated (see Fig. S2). Therefore, we limited our networks to only include individuals who were seen in at least 30% of the censuses in each of the three field expeditions in a given year (spring, summer, autumn). (Note that there were two years where censuses were only conducted during two of the three field expeditions - 1986 and 2020. In these years we retained individuals that had been seen in at least 30% of the two field expeditions).

We constructed 37 annual association networks using a ‘gambit of the group’ approach (Franks et al. 2010), whereby individuals in the same group (according to the chain rule, see above) were assumed to be associating. We estimated dyadic associations using the simple ratio index (SRI; Whitehead 2008):

SRI = Sightings_A,B_/(Sightings_A_ + Sightings_B_ − Sightings_A,B_)

where Sightings_A,B_ is the number of times individuals A and B are seen together, and Sightings_A_ and Sightings_B_ are the total number of observations of individuals A and B, respectively.

Dyadic edge weights in our network therefore range from 0 (never seen together) to 1 (never seen apart). From these annual networks, we calculated five individual level network measures designed to capture key aspects of relationship quantity (a-c) and quality (d-e). For a full breakdown of the predictions associated with each of these network metrics, see Table S1.

a. Mean group size – the average number of adult females an individual associates with per observed grouping event.
b. Degree – the number of unique adult females an individual is connected to in each annual network.
c. Participation coefficient – a measure of how well distributed an individual’s social associates are among different modules of the annual network – an individual has a high participation coefficient (i.e., 1) if it has partners uniformly distributed across all modules in the network and low participation coefficient if all partners are within its own module (i.e., 0). We characterized community structure (division of the network into distinct modules representing sub-populations that associated more with each other) in our networks using the cluster_infomap() function in igraph (Rosvall & Bergstrom 2008). We then calculated the participation coefficient using the brainGraph package (Watson 2024).
d. Mean strength – the average of an individual’s dyadic edge weights, which measures how frequently an individual associates with her partners, on average.
e. Social selectivity – the coefficient of variation in an individual’s non-zero dyadic edge weights, which measures how differentiated an individual’s social associations are. An individual with high social selectivity has strong associations with some individuals and weak associations with others, while an individual with low social selectivity has similar strength of association with all partners.

### Calculating annual home ranges

We included home range size in our analyses to account for the fact that individuals with larger home ranges were previously found to have higher annual survival (Froy et al. 2018). Following previous methodology (Froy et al. 2018; Regan et al. 2015; Wiersma et al. 2023) we estimated annual home ranges using kernel density estimation methods in the package adehabitatHR (Calenge 2006). We took the 70^th^ percent isopleth as our estimation of the core home range area (Froy et al. 2018; Regan et al. 2015; Wiersma et al. 2023).

### Estimating local density

As the density of sheep varies spatially across the study site, we anticipated that local density might influence overwinter survival above and beyond the wider effects of population density, so included it as a covariate in our models. To estimate local density we took each individual’s annual home range centroid (their mean X and Y coordinates across all recorded locations), following previous methodology (Albery et al. 2024). Using these annual centroids, we created a population-level spatial density kernel using the adehabitatHR package (Calenge 2006). This allowed us to estimate how population density is spatially distributed based on the local frequencies of individuals’ home range centroids. We then estimated an individual’s annual local density based on their centroid’s location within this spatial density kernel.

### Measuring environmental variables

We included measures of population density and winter weather in our models to quantify the extent to which social effects on survival were environmentally dependent. To measure variation in population density, we used the number of individuals in the study area recorded across the ten censuses at the end of the autumn field expedition (i.e., late November/early December). To characterize variation in the severity of winter weather conditions we used the North Atlantic Oscillation (NAO). The NAO is calculated as the surface sea-level pressure difference between Reykjavik (Iceland) and the Azores (Portugal). High NAO values are associated with warmer, windier and wetter winters across northern Europe, while low NAO values are associated with colder, drier winters (Forchhammer et al. 2002). The influences of the NAO are particularly prevalent in the winter months (December – March; (Hurrell et al. 2003)) and are known to significantly affect the Soay sheep with the increased rainfall and high winds associated with positive NAO phases leading to increased mortality (Catchpole et al. 2000; Coulson et al. 2001; Simmonds & Coulson 2015). We obtained monthly NAO data from the National Centers for Environmental Information (National Oceanic and Atmospheric Administration; https://www.ncei.noaa.gov/access/monitoring/nao/) and calculated a mean NAO for the winter period between December and March each year.

### Statistical analysis

To investigate the extent to which the quantity and quality of social associations are associated with survival, and whether these effects are moderated by environmental variables (i.e., population density and NAO) we fitted two generalized linear mixed-effects models in a Bayesian framework. Both models were fitted with a Bernoulli error distribution (logit link). Our response variable was survival to May 1^st^ of the following year (i.e., year t+1). An individual was assigned a value of 0 if it died before May 1^st^ and a value of 1 if it survived to May 1^st^ or beyond.

In both models we included a random intercept term for individual ID and observation year to account for repeated measures. We also included individual age as a random intercept as we were interested in accounting for the effect of age in our models, but not interested in estimating the effect of age on survival directly (note that our results remained the same when including age and age^2^ as fixed effects in the model, see Fig. S3).

Our ‘quantity model’ included degree and participation coefficient as continuous fixed effects while our ‘quality model’ included social selectivity and mean strength as continuous fixed effects. Although we had explicit predictions for how group size (an indirect measure of relationship ‘quantity’) might influence survival under different environmental conditions, we included group size in both models as it is expected to inherently influence social network metrics by determining the range of possible values that metrics can take (Anderson et al. 1999; De Moor et al. 2024). We included population density and NAO as continuous predictors and an interaction between each of these environmental variables and each of the included social measures in both the ‘quantity model’ and the ‘quality model’. Finally, we included home range size and local density as covariates in both models, as well as an individual’s total number of census observations, to account for any residual effects of sampling effort on social network metrics. We scaled all social metrics, as well as home range size and local density, within-year to a mean of zero and standard deviation of one, as we were interested in how relative, rather than absolute, social metrics influenced survival. We scaled population density and NAO across all years to facilitate model convergence and to allow for comparison of effect sizes between metrics.

We conducted all analyses using R version 4.1.0 (R. Core Team 2025) and fitted all models in the Bayesian software STAN (Stan Development Team 2025) using the brms package (version 2.22; Bürkner 2017). All fixed effects were given weakly informative priors. We ran all models for 20,000 iterations across four chains with a warm-up period of 2,000 iterations and thinning interval of 2. We assessed model convergence by examining trace plots to assess sampling mixing and by ensuring Rhat = 1. Statistical inference was based on the full model. We considered estimates of fixed effects to be significantly different from zero when the 95% credible intervals of the posterior distribution did not overlap zero.

## Results

On average, there were 119.74 females in each association network (range: 26-179 females) and individuals had a mean female group size of 3.99 (range: 1-9.87). Mean values of social network measures were: degree: 26 (range: 0-84); participation coefficient: 0.48 (range: 0-0.83); mean strength: 0.07 (range: 0-0.46); selectivity: 0.80 (range: 0-2.10). All social measures were weakly to moderately correlated (Fig. S4).

As expected, we found that NAO had a negative effect on annual survival, meaning that female sheep were more likely to die in wet and windy winters (quantity model: β = - 0.58; 95% CI = [-0.90, -0.26]; Fig 1A, 2A). Population density also had the anticipated negative effect on survival (quantity model: β = -0.98; 95% CI = [-1.32, -0.66]; Fig 1A, 2A). In line with previous studies, we found that home range size had a positive effect on survival (quantity model: β = 0.21; 95% CI = [0.07, 0.35]; Fig 1A, 2A). Local density also had a positive effect on survival (quantity model: β = 0.25; 95% CI = [0.11, 0.38]; Fig 1A, 2A), which likely reflects sheep congregating in areas of high-quality resources. Some measures of both relationship quantity and quality positively influenced annual survival and these effects were on par with, and sometimes stronger than the effects of home range size and local density (Fig 1A, 2A, see below).

**Figure 1.**
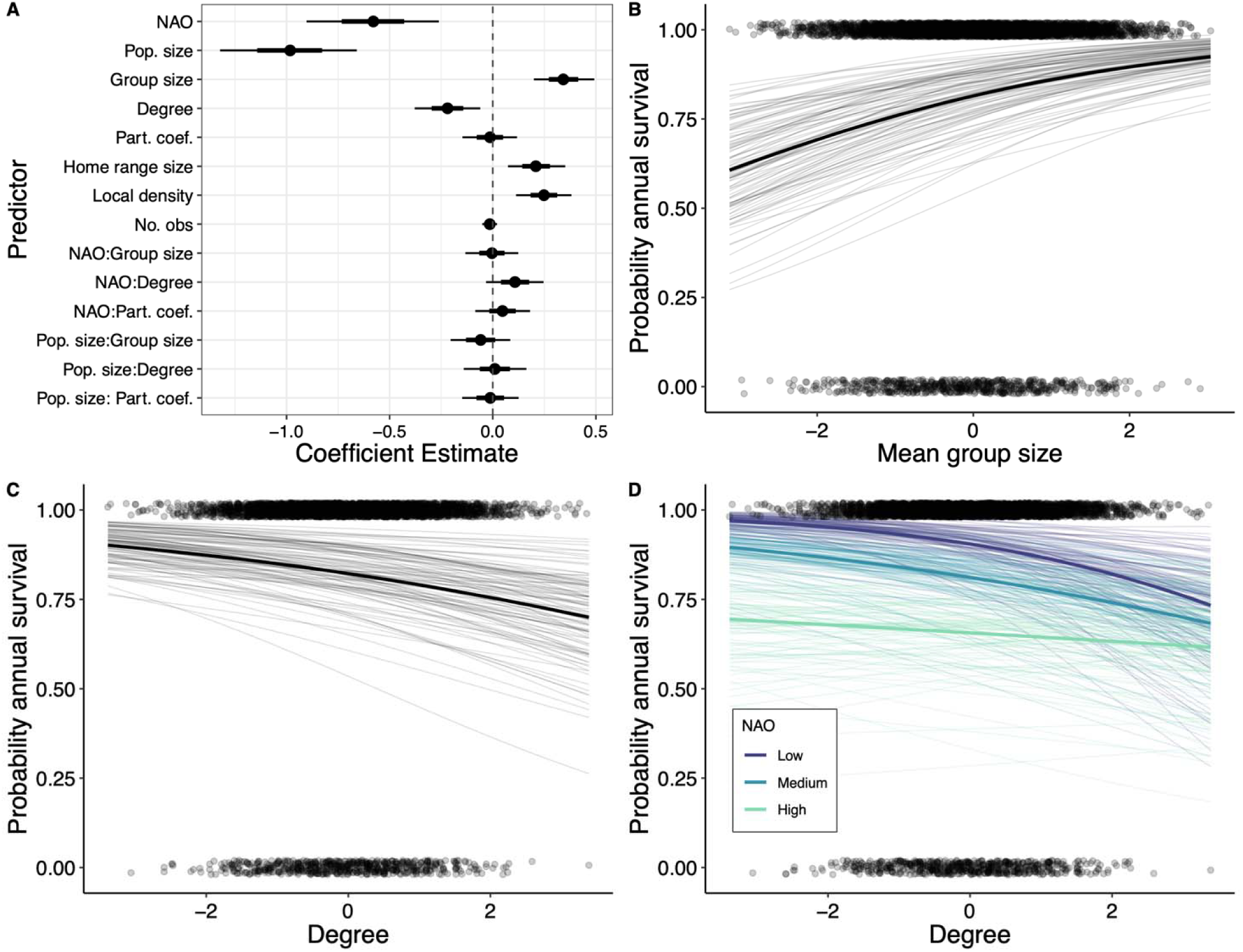
Results from the social ‘quantity’ model. (A) Parameter estimates (mean of the posterior distribution) with whiskers representing the 66% (thicker black line) and 95% (thinner black line) credible intervals (CI) for all fixed effects. (B-D) Predicted effects of (B) mean group size, (C) degree, and (D) the interaction between NAO and degree on survival. Bold lines show the mean effect with thinner lines illustrating 100 random draws of the posterior distribution. Points represent raw data. For the purposes of plotting, NAO was categorized as follows: low (1-2 SD below the mean), medium (± 0.5 SD around the mean), high (1-2 SD above the mean).

**Figure 2.**
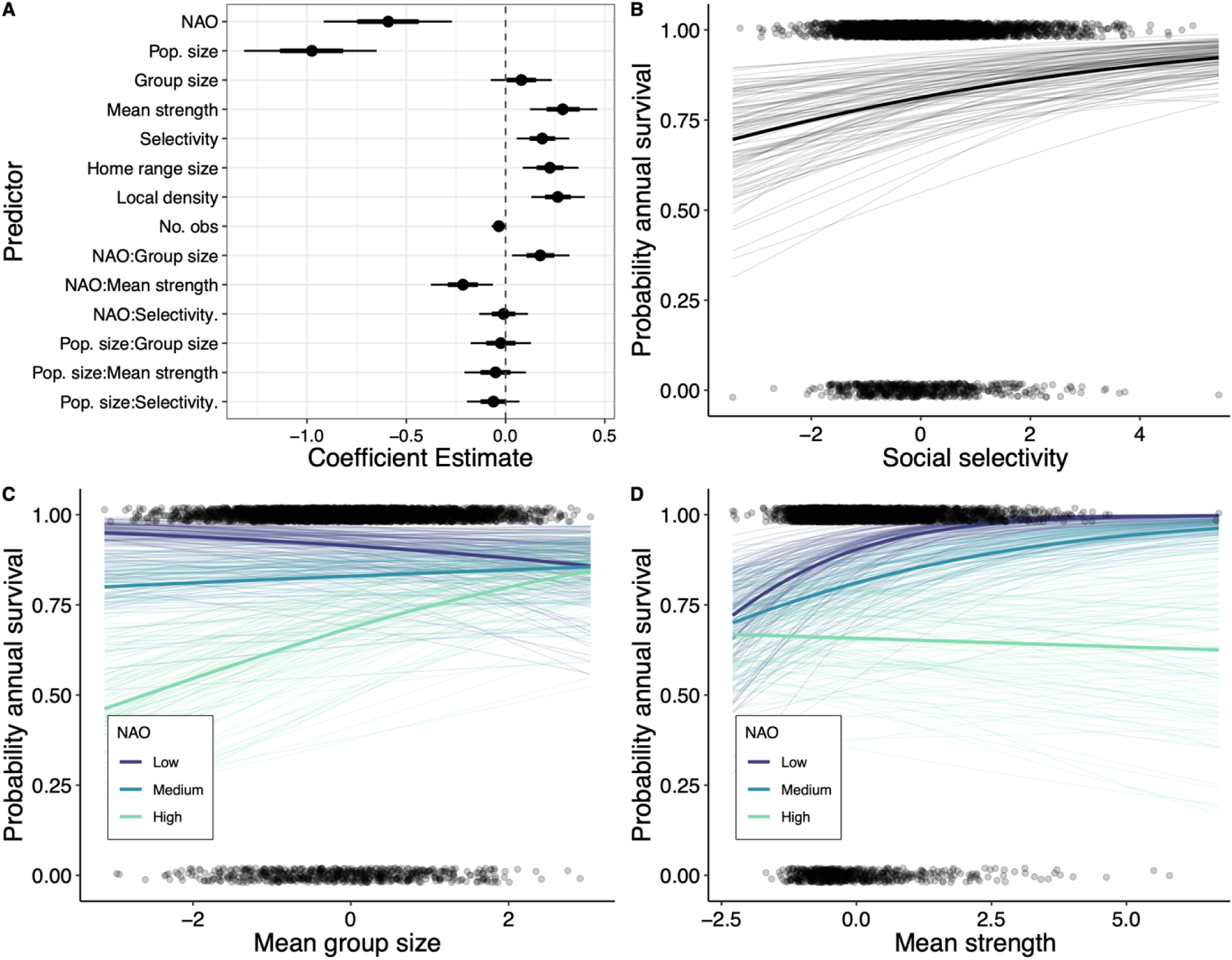
Results from the social ‘quality’ model. (A) Parameter estimates (mean of the posterior distribution) with whiskers representing the 66% (thicker black line) and 95% (thinner black line) credible intervals (CI) for all fixed effects. (B-D) Predicted effects of (B) social selectivity, (C) the interaction between NAO and mean group size, and (D) the interaction between NAO and mean strength on survival. Bold lines show the mean effect with thinner lines illustrating 100 draws of the posterior distribution. Points represent raw data. For the purposes of plotting, NAO was categorized as follows: low (1-2 SD below the mean), medium (± 0.5 SD around the mean), high (1-2 SD above the mean).

Regarding relationship ‘quantity’, the effect of mean female group size on survival was positive, although model dependent. In the ‘quantity model’ we found that survival increased with increasing group size independently of NAO (β = 0.34; 95% CI = [0.20, 0.49]; Fig 1A,B), whereas in the ‘quality model’ the positive effect of group size on survival was moderated by NAO (β = 0.18; 95% CI = [0.03, 0.32]; Fig. 3A,C). When the NAO index was high (1.5 SD above mean; wet and windy conditions), an increase in group size from 2.9 (1 SD below mean) to 5.0 (1 SD above mean) in the ‘quality model’ resulted in a 15% increase, on average, in the probability of annual survival (range: 6%-24%). Meanwhile, when the NAO index was low (1.5 SD below mean; dry and cold conditions), mean group size had no clear effect on survival in the ‘quality model’. In other words, individuals in larger groups had better survival than individuals in smaller groups in poor winter conditions, and similar survival when conditions were more benign. We did not find any evidence that the effect of group size on survival was dependent on population density in either the ‘quantity’ or the ‘quality’ model (Table 1, 2; Fig. 1A, 2A).

**Figure 3.**
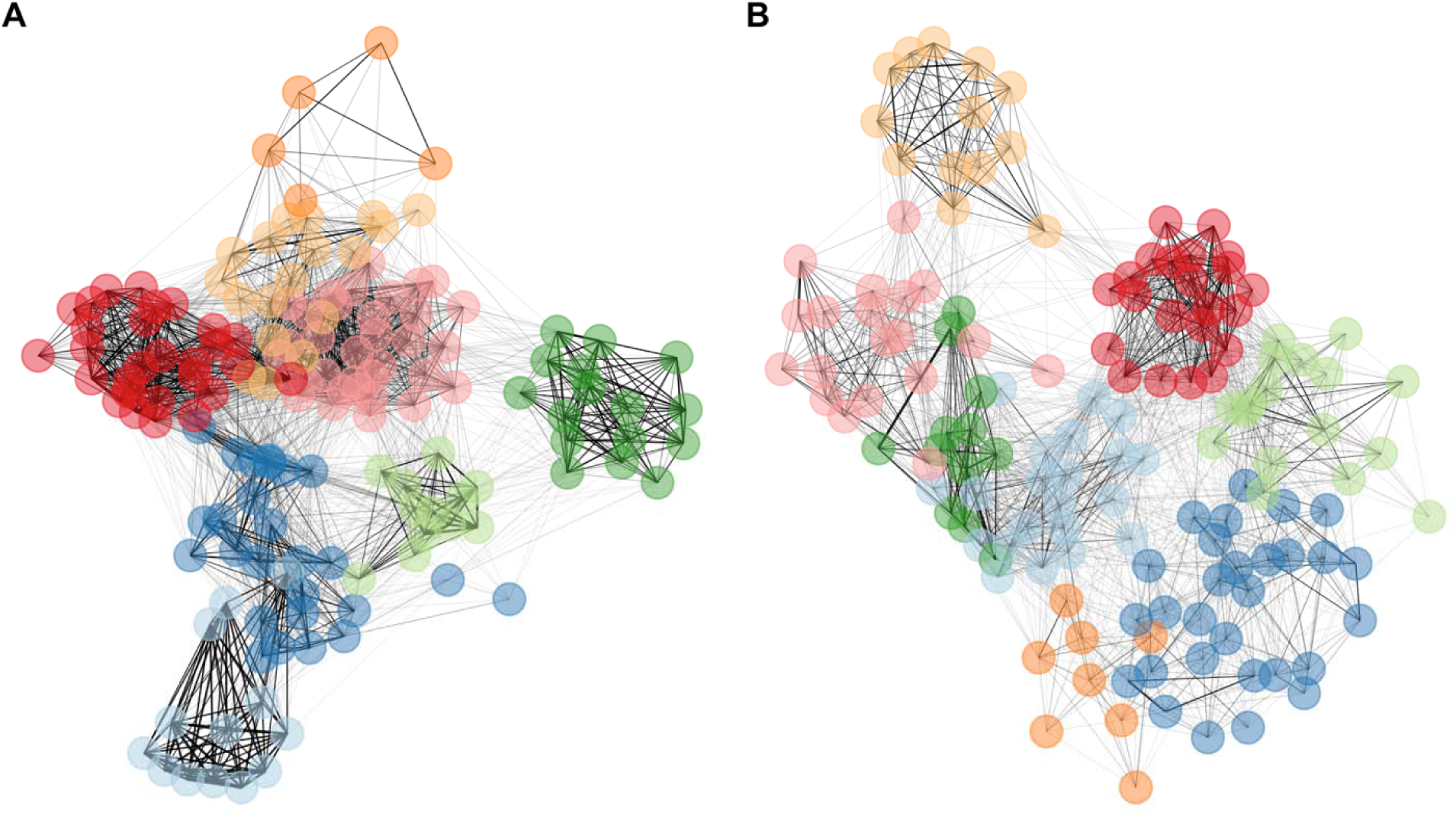
Social networks from (A) 1998, a year with a high NAO index (0.54) and (B) 2009, a year with a low NAO index (-1.48). Both years had similar population density, 591 and 617, respectively. Different colours illustrate community membership of individuals in the network.

**Table 1.** Fixed and random effects from the social ‘quantity’ model. Bolded terms indicate fixed effects where the 95% credible intervals did not overlap zero.

| Effect | Group | Term | Estimate | Lower 95% CI | Upper 95% CI |
| --- | --- | --- | --- | --- | --- |
| **Fixed Effects** |  | Intercept | 1.96 | 0.55 | 3.31 |
|  |  | **NAO** | **-0.58** | **-0.90** | **-0.26** |
|  |  | **Population size** | **-0.98** | **-1.32** | **-0.66** |
|  |  | **Group size** | **0.34** | **0.20** | **0.49** |
|  |  | **Degree** | **-0.22** | **-0.38** | **-0.06** |
|  |  | Participation coef. | -0.01 | -0.15 | 0.12 |
|  |  | **Home range size** | **0.21** | **0.07** | **0.35** |
|  |  | **Local density** | **0.25** | **0.11** | **0.38** |
|  |  | No. census obs. | -0.01 | -0.05 | 0.02 |
|  |  | NAO:Group size | 0.00 | -0.13 | 0.12 |
|  |  | NAO:Degree | 0.11 | -0.03 | 0.25 |
|  |  | NAO:Part. coef. | 0.05 | -0.08 | 0.18 |
|  |  | Pop. size:Group size | -0.06 | -0.20 | 0.08 |
|  |  | Pop. size:Degree | 0.01 | -0.14 | 0.16 |
|  |  | Pop. size:Part. coef. | -0.01 | -0.15 | 0.13 |
| **Random Effects** | Age | sd(intercept) | 1.67 | 1.02 | 2.76 |
|  | Individual.ID | sd(intercept) | 0.32 | 0.01 | 0.94 |
|  | Year | sd(intercept) | 0.84 | 0.61 | 1.16 |

In contrast, degree had an overall negative effect on survival (β = -0.22; 95% CI = [- 0.38, -0.06]; Fig. 1A,C), meaning that, conditional on mean group size, individuals with more partners had worse survival outcomes. There was some weak evidence (note that 95% credible interval overlaps 0; (Held & Ott 2018)) that the effect of degree was moderated by NAO (β = 0.11; 95% CI = [-0.03, 0.25]; Fig 1A,D), where having many partners had a negative effect on survival under conditions of low NAO (i.e., cold and dry) but no detectable effect when NAO was high (i.e., wet and windy).

We did not find any evidence that the effect of degree on survival was dependent on population density (β = 0.01; 95% CI = [-0.14, 0.16]; Fig. 1A). We also found no effect of participation coefficient on survival, either as a main effect or in interaction with either environmental variable (Table 1, Fig. 1A).

Regarding relationship ‘quality’, we found that individuals who were more selective in their social associations, that is, individuals who formed strong relationships with a few partners and weak relationships with others had higher annual survival (β = 0.19; 95% CI = [0.06, 0.32]; Fig 2A,B). An increase in social selectivity from 0.56 (1 SD below mean) to 1.01 (1 SD above mean) was associated with an average 5% increase in survival probability (range: 1%-10%). This effect of social selectivity was not moderated by either NAO or population density (Table 2; Fig 2A). We also found that individuals with high mean strength – that is, individuals who associated more frequently with the same partners – had higher survival probabilities. This effect was dependent on NAO (β = -0.22; 95% CI = [-0.36, -0.06]; Fig. 2A,D). When the NAO index was low (1.5 SD below mean; cold and dry conditions), an increase in mean strength from 0.04 (1 SD below mean) to 0.12 (1 SD above mean) resulted in a 9% increase, on average, in the probability of annual survival (range: 5%-13%). Meanwhile, when the NAO index was high (1.5 SD above mean; wet and windy conditions), mean strength had no clear effect on survival. In other words, individuals with high mean strength had better survival than individuals with low mean strength when winter conditions were benign, and similar survival when winter conditions were poor. We did not find any evidence that the effect of mean strength on survival was dependent on population density (β = -0.05; 95% CI = [0.-21, 0.10]; Fig. 2A).

**Table 2.** Fixed and random effects from the social ‘quality’ model. Bolded terms indicate fixed effects where the 95% credible intervals did not overlap zero.

| Effect | Group | Term | Estimate | Lower 95% CI | Upper 95% CI |
| --- | --- | --- | --- | --- | --- |
| **Fixed Effects** |  | Intercept | 2.45 | 1.03 | 3.83 |
|  |  | **NAO** | **-0.59** | **-0.92** | **-0.27** |
|  |  | **Population size** | **-0.98** | **-1.32** | **-0.65** |
|  |  | Group size | 0.08 | -0.07 | 0.23 |
|  |  | **Mean strength** | **0.29** | **0.12** | **0.46** |
|  |  | **Selectivity** | **0.19** | **0.06** | **0.32** |
|  |  | **Home range size** | **0.22** | **0.09** | **0.37** |
|  |  | **Local density** | **0.26** | **0.13** | **0.40** |
|  |  | No. census obs. | -0.03 | -0.07 | 0.00 |
|  |  | **NAO:Group size** | **0.18** | **0.03** | **0.32** |
|  |  | **NAO:Mean strength** | **-0.22** | **-0.38** | **-0.06** |
|  |  | NAO:Selectivity | -0.01 | -0.13 | 0.11 |
|  |  | Pop. size:Group size | -0.02 | -0.18 | 0.13 |
|  |  | Pop. size:Mean strength | -0.05 | -0.21 | 0.10 |
|  |  | Pop. size:Selectivity | -0.06 | -0.19 | 0.07 |
| **Random Effects** | Age | sd(intercept) | 1.68 | 1.03 | 2.76 |
|  | Individual.ID | sd(intercept) | 0.33 | 0.01 | 0.93 |
|  | Year | sd(intercept) | 0.86 | 0.62 | 1.18 |

## Discussion

In this paper we investigate the environment-dependent benefits of social associations in a fission-fusion social system. Our results demonstrate that being socially selective has a consistently strong, positive relationship with overwinter survival probability in adult female Soay sheep, illustrating that even in fission-fusion societies, forming strong, differentiated relationships with a few partners can carry important fitness benefits. Our findings also illustrate that the association between social network position and survival in adult female sheep depends on weather conditions but not population density. In general, we find that having many social associations helps to buffer individuals against harsh overwinter conditions, in line with our predictions, while maintaining consistent and strong social relationships is more important under benign conditions. Together, this provides some of the first evidence that the fitness benefits of sociality are environment-dependent.

Our findings contribute to the growing evidence highlighting the importance of an individual’s social network for its fitness in non-human animals (Formica et al. 2012; McDonald 2007; Riehl & Strong 2018; Snyder-Mackler et al. 2020). In particular, our results help improve understanding of the benefits of specific social association patterns in fission-fusion systems, which remain understudied. The potential benefits of gregariousness, especially in socially flexible ungulates, is well-established (Kie 1999), and we offer further support for this idea by showing that mean group size has a broadly positive effect on survival. However, the importance of forming strong, stable relationships, while generally accepted to be important in species living in stable social groups, remains less certain in these more fluid social systems (but see Connor & Krützen 2015). Previous studies have demonstrated positive associations between an individual’s social centrality and survival (Stanton & Mann 2012; Vander Wal et al. 2015) or reproductive success (Feldblum et al. 2021) this pattern isn’t universal (Montero et al. 2020). Our findings show that maintaining strong, consistent social associations with a subset of potential partners positively predicts survival, thus demonstrating that developing differentiated relationships with a smaller number of individuals can be important in fission-fusion societies. The most likely explanation for this finding is that keeping a familiar social environment is beneficial to individuals, perhaps by enhancing the predictability of social interactions and thus stabilising dominance interactions (Robinson & Kruuk 2007), reducing the potential for conflict and aggression and increasing foraging efficiency (Fox et al. 2024; Griffiths et al. 2004; Siracusa et al. 2019; Weidt et al. 2008). However, further empirical and theoretical work is required to investigate the mechanisms that underlie this relationship.

Investigating how different aspects of an individual’s social network vary in how they shape fitness offer an important window into the evolutionary function of these relationships. While previous studies have typically linked fitness-related traits, such as survival, to a single aspect of an individual’s social network position, there is increasing recognition that there are different ways of being socially connected that might matter in different contexts (Cheney et al. 2016; Ellis et al. 2019; McFarland et al. 2017; Sabol et al. 2020; Schülke et al. 2022; De Moor et al. preprint). Overall, our results show that relationship ‘quality’ strongly predicts adult female overwinter survival, with a persistent positive effect of social selectivity and environment-dependent positive effect of mean association strength on survival. In contrast, the ‘quantity’ of an individuals’ social associations has a more mixed effect. While being in larger groups is generally associated with improved survival, having more partners is not, and, in fact, has a negative effect once group size is accounted for. Importantly, we show that the relative importance of relationship ‘quality’ and ‘quantity’ is context-dependent. Specifically, we demonstrate a clear interaction between the effects of winter weather and social network position on overwinter survival of adult female Soay sheep. During windy, wet winters, winter survival is positively associated with group size, whereas during more benign winters, the strength of an individual’s connections to its social partners better predicts survival. In addition, having more partners has the strongest negative effect on survival during benign winters, which are the same conditions under which stronger connections to partners are favoured. Taken together, our findings join a growing body of literature that suggests that the fitness benefits of sociality are context dependent (Ellis et al. 2017; Fisher & Cheney 2023), with more benign conditions favouring ‘quality’ and harsher conditions favouring ‘quantity’ in our study system. Determining the causes of this complex environmental dependency will demand further in-depth behavioural and demographic study. As outlined above, we suspect that familiarity with social partners is important, reducing competition and conflict for individuals with high relationship quality. Larger group size, on the other hand, might provide thermoregulatory benefits, with particular benefit to survival during harsh winters when these thermoregulatory benefits are most important. Indeed, during storms the Soay sheep are often observed huddling in the shelter provided by walls and small stone buildings (‘cleits’), and it seems highly feasible that larger groups may enhance the benefits of such huddling/sheltering behaviours.

Our results also point toward a trade-off between quality and quantity – where individuals can either have many social associates or interact consistently with the same few partners. Such trade-offs have been suggested to occur in other species with more stable social systems and may be a consequence of energetic or time constraints (Dakin & Ryder 2020; Dunbar 2020; Korstjens et al. 2006; Lehmann et al. 2007; de Silva et al. 2011). In fission-fusion social systems, like the Soay sheep, an individual’s social connectedness depends on both its gregariousness (group size preference) and its tendency to remain with the same social partners or move between potential partners. Both of these likely vary consistently among individuals (Godde et al. 2013). In our results degree and mean strength are positively correlated with an individual’s mean group size across a year. More gregarious individuals tend to have more partners and, likely because they are located more centrally within the population, tend to interact more frequently with the same partners. However, once group size is accounted for, we find that degree and mean strength are strongly negatively correlated (Pearson’s r = -0.8). This suggests a trade-off: females either consistently group with the same partners and remain well-integrated in their group, resulting in high relationship strength with those few partners, or “float” and maintain less consistent group composition, resulting in many social partners, but lower strength with those partners. Our results indicate that it is these “floater females” who suffer poorer overwinter survival. Given this pattern for “floater females”, it is perhaps surprising that high participation coefficients (i.e., social associates widely dispersed among network “communities”) are not associated with reduced survival, but this can be explained if social communities cover broader spatial areas than each individual’s typical home range.

We also had strong expectations that the relationship between social network position and survival would depend on population density, in particular through its effects on access to food. However, we did not find evidence that the effects of any of our social network measures on adult female survival were moderated by global population density, despite global density having the expected negative effect on survival (Coulson et al. 2001). The lack of an interactive effect between population density and sociality on survival may depend on the proposed mechanism linking social associations to foraging opportunities. Given the widespread nature of food availability for an herbivore, social relationships may be less important in governing access to food than they might otherwise be in species where resources are more centralized and defendable (Jungwirth et al. 2021). However, familiarity with conspecifics might still facilitate a feeling of ‘safety’ when foraging in groups by reducing generalized aggression and thus social vigilance (Favreau et al. 2015; Gaynor & Cords 2012; Josephs et al. 2016; Szulanski et al. 2023), thereby enhancing foraging time and/or efficiency (Griffiths et al. 2004). This role of familiarity might help to explain why maintaining strong and consistent relationships generally benefit survival but why these relationships are no more important in years of high competition when access to food is scarce.

A key limitation of our study is that we only had association data from which to derive metrics of social connectedness. Given that association data only represent a proxy for behavioural interactions (Weiss et al. 2021), it is impossible to disentangle whether the patterns of social connectedness we observe are driven by social preferences per se, or emerge from fine-scale patterns of spatial behaviour (Webber et al. 2023), which limits the strength of our conclusions. We were careful in our statistical models to control for factors such as local density and home range area that may be correlated with measures of social connectedness, and also have a direct effect on overwinter survival or survival-associated traits (Albery et al. 2024; Froy et al. 2018). However, it is possible that social network measures vary across space (see Albery et al. 2021) for reasons that are harder to control for directly (e.g., habitat configuration), preventing us from being wholly confident of a causal relationship between social connectedness and survival. Future research would benefit from the collection of behavioural interaction data in this system to help disentangle spatial and social behaviour. Collecting such data over shorter timescales and pairing it with longer term patterns in association data would allow us to identify the mechanisms by which social associations impact individual fitness. We also flag that we are using measures of social connectedness from the spring to autumn when most mortality occurs in winter. Given our results, we feel it is likely that individuals display relatively consistent social phenotypes (as per Aplin et al. 2015; Blaszczyk 2018; Evans et al. 2021; Strickland et al. 2021) and that social behaviour during the summer may also contribute to overwinter survival through carry-over effects (Harrison et al. 2011). Using bio-logging devices to enable social data collection across the entire year could investigate the relative importance of these two explanations.

In our study we have demonstrated strong relationships between individual social network position and overwinter survival for adult female Soay sheep. In particular, our findings highlight the likely importance of maintaining strong associations with a subset of potential social partners for a species with fluid fission-fusion social dynamics, indicating that the importance of maintaining ‘quality’ social relationships extends beyond species that live in stable social systems. However, we also demonstrate that the relative importance of the ‘quality’ and ‘quantity’ of social relationships is partially dependent on environmental conditions, potentially providing a key mechanism by which individual variation in social behaviour can be maintained. As such, future work should also consider how context-dependence of social benefits is linked with phenotypic plasticity as a further explanation for this variation. We note that it is only by attempting to break down the different aspects of an individual’s ‘social connectedness’ that we could gain a fuller understanding of these relationships between social behaviour and survival and the mechanisms that might underlie them. As such, we encourage researchers to use similar, multifaceted approaches in future. Collectively, our results help generalize our understanding of the evolutionary ecology of social networks in wild populations, pointing towards the value of considering temporal dynamics of social behaviour and its consequences across diverse social systems in future theoretical and empirical research.

## Acknowledgements

We thank all the personnel who have been involved in the long-term study of Soay sheep on St Kilda, Scotland, over the last 39 years, particularly Ian Stevenson, Michael Morrisey, Jon Slate, Susan Johnston, Tim Clutton-Brock, Loeske Kruuk and the many field volunteers. We are also grateful to the National Trust for Scotland for permission to work on St Kilda, and QinetiQ and Kilda Cruises for logistical assistance in the field.

M.J.S. is supported by a Royal Society University Research Fellowship URF\R1\2 21 800. D.D.M. is funded by the Max Planck Society.

