## Supplementary Materials for "Environment dependent benefits of sociality in Soay sheep"

**This PDF file includes:**

Table S1
Figures S1 to S4

As noted in the main manuscript, individual female sheep were typically only recorded the first time they were seen on a given census route, even though they might have been observed again in another group later in the census. For simulations showing that this is unlikely to substantially affect network structure or conclusions drawn about individual social network position, please see the attached code labeled “BiasMarkdown.html” and data labeled “sheep_year_example.csv”.

**Table S1**. Predicted effects of different measures of social association quantity (green) or quality (purple) on female survival under different environmental conditions.

| **Social network metric** | **NAO prediction**  *(high = wet & windy)* | **NAO rationale** | **Population density prediction** | **Population density rationale** |
| --- | --- | --- | --- | --- |
| **Mean group size**  *(average number of females per group)* | Larger mean group size predicted to **increase** survival when NAO high | Larger groups will help shelter individuals from adverse conditions and provide thermoregulatory benefits. | Larger mean group size predicted to **reduce** survival when density high | Individuals in larger groups will suffer greater competition when resources are limited. |
| **Degree**  *(number of unique partners)* | Higher degree predicted to **increase** survival when NAO high | Having more social associates will improve social tolerance, improving access to more groups or cleits and to more central locations within groups or cleits. | Higher degree predicted to **increase** survival when density high | Having more social associates will improve social tolerance around limited resources and social knowledge of where high-quality resources are located. |
| **Participation coefficient** *(PC; distribution of partners among modules in the network)* | Exact prediction depends on underlying mechanism *(inclusion allows us to disentangle how degree influences survival)* | If benefits come from access to many groups or cleits, then widespread social associations across many modules in the population ( *high PC)* will enhance survival when NAO high   If benefits come from accessing preferred cleits or central locations within groups, then having many social associations within your module ( *low PC*) will enhance survival when NAO high | Exact prediction  depends on underlying mechanism *(inclusion allows us to disentangle how degree influences survival)* | If benefits come from access to many resource patches or knowledge of where resources are, then widespread social associations across many modules in the population (*high PC*) will enhance survival when density high  If benefits come from improved social tolerance around limited resources, then many social associations within your module (*low PC*) will enhance survival when density high |
| **Mean strength** *(average frequency of association with partners)* | Effect on survival **not expected** to depend on NAO | Strong overall connection to associates and strongly differentiated relationships are not expected to be important for accessing cleits | Higher mean strength predicted to **increase** survival when density high | Strong overall connection to associates will improve social tolerance |
| **Social selectivity** *(differentiation in frequency of association with partners)* | Effect on survival **not expected** to depend on NAO |  | Effect on survival **not expected** to depend on density | Strongly differentiated relationships not expected to be important for accessing food resources |

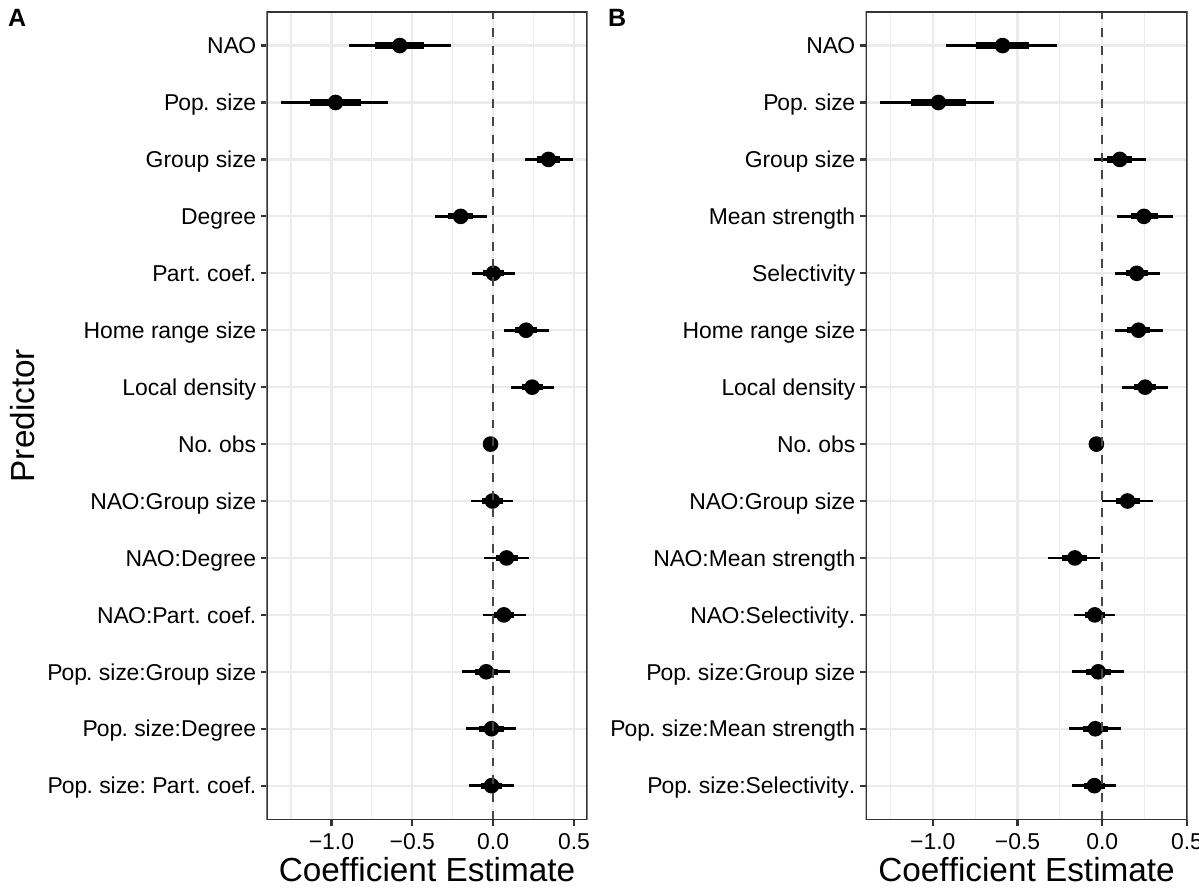

**Figure S1.** Results from the (A) social ‘quantity’ model and (B) social ‘quality’ model fitted when retaining only the first observation for each female sheep from a given census. Parameter estimates (mean of the posterior distribution) are shown with whiskers illustrating that 66% and 95% credible intervals (CI) for all fixed effects.

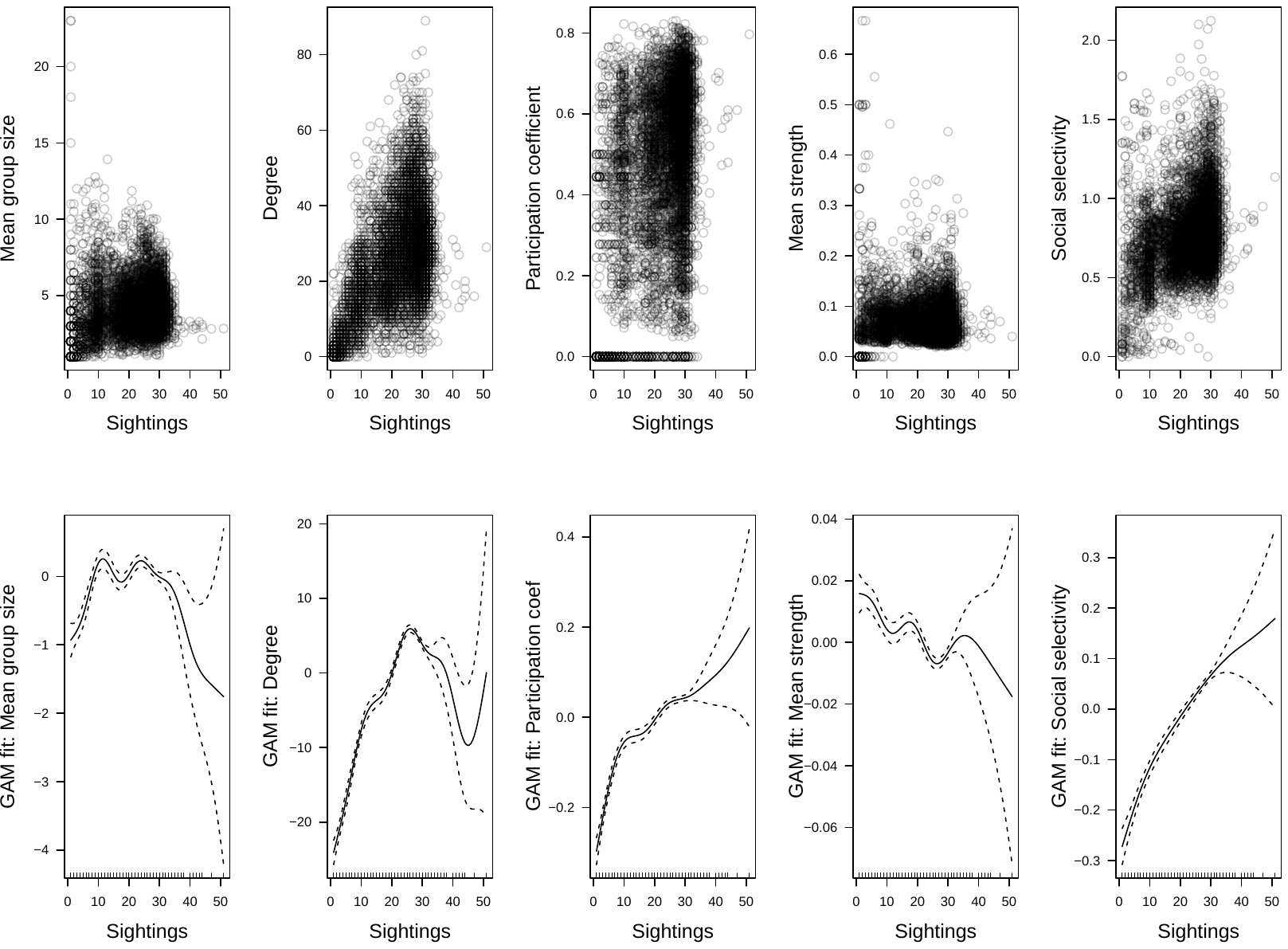

**Figure S2.** Effect of number of census observations on social network measures. Top panels show plots of raw data, bottom panels show a GAM smoothing function fit through raw data.

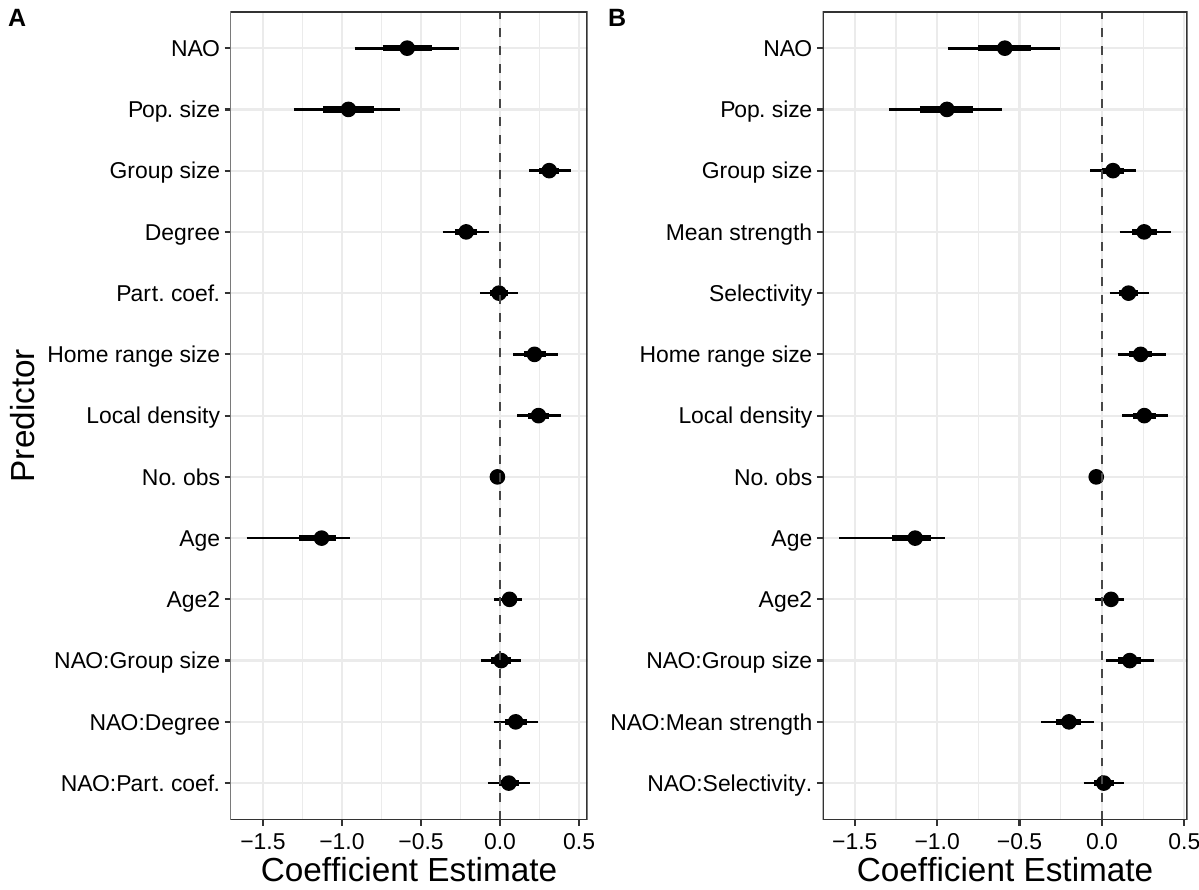

**Figure S3.** Results from the (A) social ‘quantity’ model and (B) social ‘quality’ model fitted when including age and age^2^ as fixed effects. Parameter estimates (mean of the posterior distribution) are shown with whiskers illustrating that 66% and 95% credible intervals (CI) for all fixed effects.

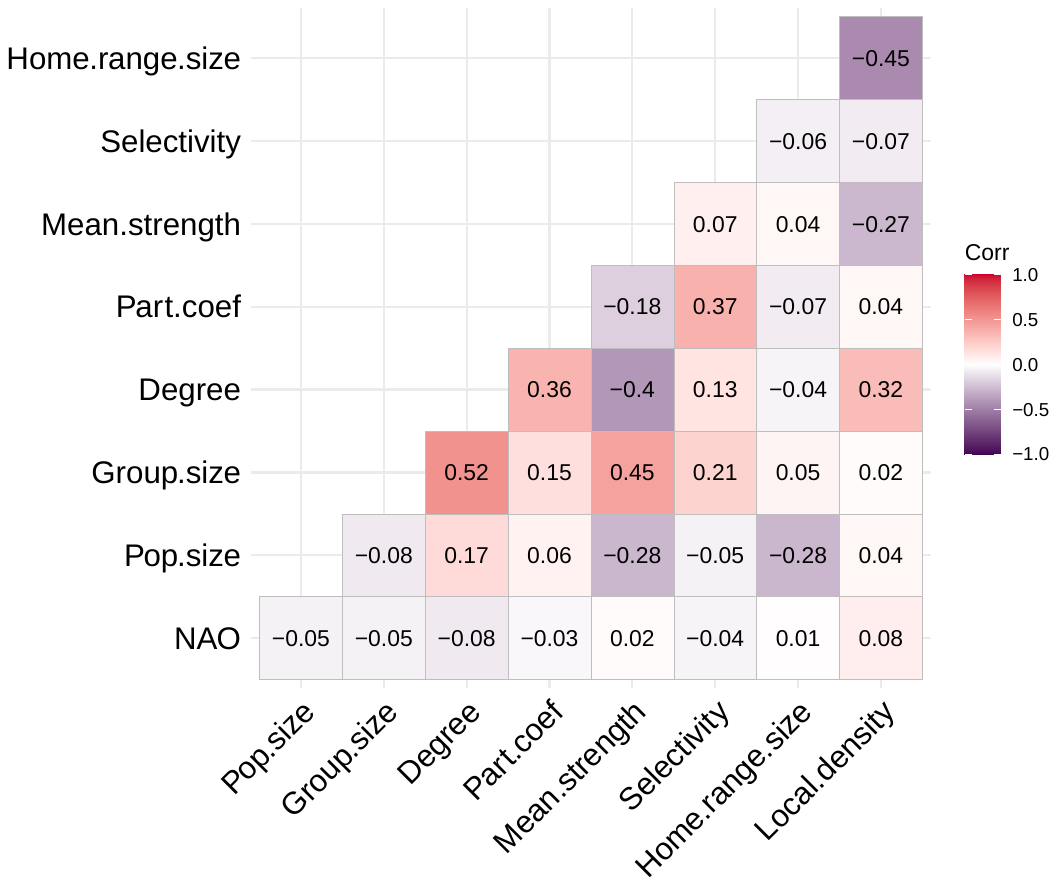

**Figure S4.** Pearson correlation coefficients (r) between all model covariates.
