## Supplementary Code for "Environment dependent benefits of sociality in Soay sheep"

Soay Sheep networks - testing for bias caused by the census approach


### Soay Sheep networks - testing for bias caused by the census approach

###### Matthew Silk

#### 2025-04-15

##### Load required R packages

Load the R packages we will use in the analysis

```
library(igraph)
```

```
## Warning: package 'igraph' was built under R version 4.1.3
```

```
## 
## Attaching package: 'igraph'
```

```
## The following objects are masked from 'package:stats':
## 
##     decompose, spectrum
```

```
## The following object is masked from 'package:base':
## 
##     union
```

```
library(asnipe)
```

##### Load data

We now load the example network we will use to test how sampling
affects measured propertes of the network

```
path<-"C:/Users/matth/Dropbox/Edinburgh/SoaySheep/BasicNetworkExample"
data2<-read.csv(paste0(path,'/sheep_year_example.csv'))
```

##### Create group-by-individual matrix

In this code chunk we generate a group-by-individual matrix with the
cesnsus data from our chosen year

```
#List of unique ideas and the number of sheep
ids2<-unique(data2$ID)
ids2<-ids2[!is.na(ids2)]
n.sheep2<-length(ids2)

#Empty matrix for the network
network_full<-matrix(0,nr=n.sheep2,nc=n.sheep2)
rownames(network_full)<-colnames(network_full)<-ids2

#New date as factor to be easier to work with
data2$date_fac<-as.numeric(as.factor(data2$Date))

#New column to create unique combos of date and group
data2$groupno2<-paste0(data2$date_fac,"_",data2$GroupNo)

#Unique groups and number of groups
groups2<-unique(data2$groupno2)
n.groups2<-length(groups2)

#Creates a group-by-individual matrix and fills it
#Groups are rows, columns are individual. 1 indicates an individual in a group
gbi<-matrix(0,nr=n.groups2,nc=n.sheep2)
for(i in 1:nrow(data2)){
  if(is.na(data2$ID[i])==FALSE){
    gbi[which(groups2==data2$groupno2[i]),which(ids2==data2$ID[i])]<-1
  }
}
```

##### Inspect social network

Here we will quickly check that the code has generated a sensible
social network

```
net1<-asnipe::get_network(gbi)
```

```
## Generating  449  x  449  matrix
```

```
net1_i<-igraph::graph_from_adjacency_matrix(net1,diag=NA,mode="undirected",weighted=TRUE)
plot(net1_i,vertex.label=NA,vertex.size=4)
```

Now we have generated a social network from the data we have
observed

---


---

#### Simulation 1: Missing observations are random

Next is to create a potential scenario which includes the missed
observations that may occur. These are observations of the same sheep on
the same census route if it has moved between groups. Consequently our
approach is to sample a particular sheep and then add it at random to
any group recorded later on the same route.

To start with we do this by selecting a sheep at random and adding it
to a random group (this is likely conservative relative to our observed
data in terms of the noise added to the data).

Here we test missing 15 observation of missing sheep (a conservative
estimate of the expected rate of this occurring provided by the field
teams). However, to ensure the outcome is robust to this choice of
threshold we have repeated this with arbitrarily high thresholds up to
60 (much higher than that expected to occur) and our results remain
robust (even if some correlations weaken slightly) well beyond the level
at which these missing observation occur in the study.

##### Create unobserved ‘realised’ network

Here we start by selecting sheep at random and use a threshold of
15.

```
#Replicate census data
data2_new2<-data2

#Set threshold
thresh<-15

#Add sheep to the census data
censuses<-unique(data2$date_fac)
for (i in 1:length(censuses)){
  counter<-1
  data_tmp<-data2_new2[data2_new2$date_fac==censuses[i],]
  while(counter<(thresh+1)){
    
    #Select potential sheep
    sta_row<-sample(1:nrow(data_tmp),1,replace=FALSE)
    sta_gr<-data_tmp$GroupNo[sta_row]
    sta_rte<-ifelse(sta_gr<101,1,ifelse(sta_gr<201,2,3))
    
    #Adds selected sheep to new group
    pot_gr<-unique(data_tmp$GroupNo)
    if(sta_rte==1){
      pot_gr2<-pot_gr[pot_gr>sta_gr&pot_gr<101]
    }
    if(sta_rte==2){
      pot_gr2<-pot_gr[pot_gr>sta_gr&pot_gr<201]
    }
    if(sta_rte==3){
      pot_gr2<-pot_gr[pot_gr>sta_gr&pot_gr>200]
    }
    
    if(length(pot_gr2)>1){
      new_gr<-sample(pot_gr2,1,replace=FALSE)
    }
    if(length(pot_gr2)==1){
      new_gr<-pot_gr2
    }
    if(length(pot_gr2)==0){
      new_gr<-numeric()
    }
    
    #Adjusts corresponding census data
    if(length(new_gr)>0){  
      if(new_gr!=sta_gr&sum(data_tmp$ID[data_tmp$GroupNo==new_gr]%in%data_tmp$ID[sta_row])==0){
        data_tmp2<-data_tmp[data_tmp$GroupNo==new_gr,]
        new_data<-data_tmp2[1,]
        new_data$ID<-data_tmp$ID[sta_row]
        new_data$Sex<-data_tmp$Sex[sta_row]
        new_data$Age<-data_tmp$Age[sta_row]
        data_tmp<-rbind(data_tmp,new_data)
        counter<-counter+1
      }
    }
    
  }
  
  #Adds new data to census dataset
  data2_new2<-data2_new2[!data2_new2$date_fac==censuses[i],]
  data2_new2<-rbind(data2_new2,data_tmp)
  
}

#List of unique ideas and the number of sheep
ids2<-unique(data2_new2$ID)
ids2<-ids2[!is.na(ids2)]
n.sheep2<-length(ids2)

#Unique groups and number of groups
groups2<-unique(data2_new2$groupno2)
n.groups2<-length(groups2)

#Creates a group-by-individual matrix and fills it
#Groups are rows, columns are individual. 1 indicates an individual in a group
gbi<-matrix(0,nr=n.groups2,nc=n.sheep2)
for(i in 1:nrow(data2_new2)){
  if(is.na(data2_new2$ID[i])==FALSE){
    gbi[which(groups2==data2_new2$groupno2[i]),which(ids2==data2_new2$ID[i])]<-1
  }
}

#Create adjacency matrix of new realised network
net3<-asnipe::get_network(gbi)
```

```
## Generating  449  x  449  matrix
```

```
#Create corresponding network object
net3_i<-igraph::graph_from_adjacency_matrix(net3,diag=NA,mode="undirected",weighted=TRUE)
```

We can now test how this affects the network.

##### How are the two networks correlated?

Just a simple plot of one network against the other.

```
plot(net1~net3,pch=16,col=adjustcolor("black",0.3),xlab="Observed network",ylab="Realised network")
```

There is a strong correlation between the two.

Although adding sheep (randomly) to extra groups caused lower SRIs in
general. This is unsurprising given we are adding them to random
groups.

```
sum(net3<net1)
```

```
## [1] 20152
```

```
sum(net3>net1)
```

```
## [1] 3252
```

We can then test the affect on the types of network measure used in
the study. We will focus on degree and strength as these (combined with
the raw SRI values) underlie most the measures we use.

Note here that the lines on plots are 1:1 lines and not lines of best
fit

```
#Calculate degree
deg1<-degree(net1_i)
deg2<-degree(net3_i)

#Calculate strength
str1<-strength(net1_i)
str2<-strength(net3_i)

#Plot relationship
par(mfrow=c(1,2))
par(xpd=FALSE)
plot(deg2,deg1,xlab="Degree in realised network",ylab="Degree in observed network")
lines(x=c(0,1000),y=c(0,1000))
par(xpd=FALSE)
plot(str2,str1,xlab="Strength in realised network",ylab="Strength in observed network")
lines(x=c(0,1000),y=c(0,1000))
```

```
par(mfrow=c(1,1))

#Test correlation
cor.test(deg2,deg1)
```

```
## 
##  Pearson's product-moment correlation
## 
## data:  deg2 and deg1
## t = 155.56, df = 447, p-value < 2.2e-16
## alternative hypothesis: true correlation is not equal to 0
## 95 percent confidence interval:
##  0.9890424 0.9924277
## sample estimates:
##       cor 
## 0.9908903
```

```
cor.test(str2,str1)
```

```
## 
##  Pearson's product-moment correlation
## 
## data:  str2 and str1
## t = 282.57, df = 447, p-value < 2.2e-16
## alternative hypothesis: true correlation is not equal to 0
## 95 percent confidence interval:
##  0.9966449 0.9976842
## sample estimates:
##       cor 
## 0.9972125
```

In other code we have repeated this analysis on other centrality
measures such as eigenvector centrality and betweenness centrality and
you obtain similar (but less strong) positive correlations, especially
when the threshold is very low.

---


---

#### Simulation 2: Missing observations are biased

It is feasible (although we are speculating) that sheep observed that
are missing, having been observed in multiple groups on the same route,
are a biased subset of individuals. Perhaps the most likely bias is that
individuals with higher degree centrality (i.e. more social connections)
are more likely to occur in different groups. So we repeat our
simulations but biasing our sample of sheep towards those with higher
degree.

Here we test missing 15 observation of missing sheep (a conservative
estimate of the expected rate of this occurring provided by the field
teams). However, to ensure the outcome is robust to this choice of
threshold we have repeated this with arbitrarily high thresholds up to
60 (much higher than that expected to occur) and our results remain
robust (even if some correlations weaken slightly) well beyond the level
at which these missing observation occur in the study.

```
#Replicate census data
data2_new3<-data2

#Set threshold
thresh<-15

#Add sheep to the census data
censuses<-unique(data2$date_fac)
for (i in 1:length(censuses)){
  counter<-1
  data_tmp<-data2_new3[data2_new3$date_fac==censuses[i],]
  while(counter<(thresh+1)){
    
    #Select potential sheep
    deg_tmp<-degree(net1_i)
    deg_tmp<-deg_tmp/(n.sheep2-1)
    samp_probs<-rep(NA,nrow(data_tmp))
    
    for(j in 1:length(samp_probs)){
      if(is.na(data_tmp$ID[j])==FALSE){
        samp_probs[j]<-deg_tmp[which(ids2%in%data_tmp$ID[j])]+0.001
      }
      if(is.na(data_tmp$ID[j])==TRUE){
        samp_probs[j]<-min(deg_tmp)+0.001
      }
    }
    sta_row<-sample(1:nrow(data_tmp),1,replace=FALSE,prob=samp_probs)
    sta_gr<-data_tmp$GroupNo[sta_row]
    sta_rte<-ifelse(sta_gr<101,1,ifelse(sta_gr<201,2,3))
    
    #Adds selected sheep to new group
    pot_gr<-unique(data_tmp$GroupNo)
    if(sta_rte==1){
      pot_gr2<-pot_gr[pot_gr>sta_gr&pot_gr<101]
    }
    if(sta_rte==2){
      pot_gr2<-pot_gr[pot_gr>sta_gr&pot_gr<201]
    }
    if(sta_rte==3){
      pot_gr2<-pot_gr[pot_gr>sta_gr&pot_gr>200]
    }
    
    if(length(pot_gr2)>1){
      new_gr<-sample(pot_gr2,1,replace=FALSE)
    }
    if(length(pot_gr2)==1){
      new_gr<-pot_gr2
    }
    if(length(pot_gr2)==0){
      new_gr<-numeric()
    }
    
    #Adjusts corresponding census data
    if(length(new_gr)>0){  
      if(new_gr!=sta_gr&sum(data_tmp$ID[data_tmp$GroupNo==new_gr]%in%data_tmp$ID[sta_row])==0){
        data_tmp2<-data_tmp[data_tmp$GroupNo==new_gr,]
        new_data<-data_tmp2[1,]
        new_data$ID<-data_tmp$ID[sta_row]
        new_data$Sex<-data_tmp$Sex[sta_row]
        new_data$Age<-data_tmp$Age[sta_row]
        data_tmp<-rbind(data_tmp,new_data)
        counter<-counter+1
      }
    }
    
  }
  
  #Adds new data to census dataset
  data2_new3<-data2_new3[!data2_new3$date_fac==censuses[i],]
  data2_new3<-rbind(data2_new3,data_tmp)
  
}

#List of unique ideas and the number of sheep
ids2<-unique(data2_new3$ID)
ids2<-ids2[!is.na(ids2)]
n.sheep2<-length(ids2)

#Unique groups and number of groups
groups2<-unique(data2_new3$groupno2)
n.groups2<-length(groups2)

#Creates a group-by-individual matrix and fills it
#Groups are rows, columns are individual. 1 indicates an individual in a group
gbi<-matrix(0,nr=n.groups2,nc=n.sheep2)
for(i in 1:nrow(data2_new3)){
  if(is.na(data2_new3$ID[i])==FALSE){
    gbi[which(groups2==data2_new3$groupno2[i]),which(ids2==data2_new3$ID[i])]<-1
  }
}

#Create adjacency matrix of new realised network
net4<-asnipe::get_network(gbi)
```

```
## Generating  449  x  449  matrix
```

```
#Create corresponding network object
net4_i<-igraph::graph_from_adjacency_matrix(net4,diag=NA,mode="undirected",weighted=TRUE)
```

We can again test how this affects the network.

##### How are the two networks correlated?

Just a simple plot of one network against the other.

```
plot(net1~net4,pch=16,col=adjustcolor("black",0.3),xlab="Observed network",ylab="Realised network")
```

There remains a strong correlation between the two.

The effect on bias in SRIs is now much less pronounced

```
sum(net4<net1)
```

```
## [1] 20858
```

```
sum(net4>net1)
```

```
## [1] 3990
```

We can again inspect correlations with on degree and strength. Note
again that the lines on plots are 1:1 lines and not lines of best fit.
Once again, we have checked for other (less relevant) measures and the
patterns revbealed here are qualitatively robust as would be expected
given the similarity of the networks themselves.

```
#Calculate degree
deg1<-degree(net1_i)
deg2<-degree(net4_i)

#Calculate strength
str1<-strength(net1_i)
str2<-strength(net4_i)

#Plot relationship
par(mfrow=c(1,2))
par(xpd=FALSE)
plot(deg2,deg1,xlab="Degree in realised network",ylab="Degree in observed network")
lines(x=c(0,1000),y=c(0,1000))
par(xpd=FALSE)
plot(str2,str1,xlab="Strength in realised network",ylab="Strength in observed network")
lines(x=c(0,1000),y=c(0,1000))
```

```
par(mfrow=c(1,1))

#Test correlation
cor.test(deg2,deg1)
```

```
## 
##  Pearson's product-moment correlation
## 
## data:  deg2 and deg1
## t = 147.48, df = 447, p-value < 2.2e-16
## alternative hypothesis: true correlation is not equal to 0
## 95 percent confidence interval:
##  0.9878293 0.9915877
## sample estimates:
##       cor 
## 0.9898807
```

```
cor.test(str2,str1)
```

```
## 
##  Pearson's product-moment correlation
## 
## data:  str2 and str1
## t = 272.52, df = 447, p-value < 2.2e-16
## alternative hypothesis: true correlation is not equal to 0
## 95 percent confidence interval:
##  0.9963941 0.9975110
## sample estimates:
##       cor 
## 0.9970041
```

The bias in whether individuals are missed now leads to more
pronounced underestimation (normally, note these are stochastic
simulations!) of the degree of highly connected individuals (minimal
effect on strength) but relatively speaking there remains a high
correlation such that highly connected individuals are still identified
as highly connected and vice versa.

We are confident these results demonstrate that the sampling approach
does not have a strong effect on our conclusions.

---


---


---
